# RFdiffusion Exhibits Low Success Rate in De Novo Design of Functional Protein Binders for Biochemical Detection

**DOI:** 10.1101/2025.02.07.636769

**Authors:** Bruce Jiang, Xiaoxiao Li, Amber Guo, Moris Wei, Jonny Wu

**Author notes:** Correspondence should be addressed to. Bruce Jiang.

## Abstract

The design of high-affinity protein binders is critical for biochemical detection, yet traditional methods remain labor-intensive. AI-driven tools like RFdiffusion, a RoseTTAFold-based diffusion model, offer promising alternatives for generating protein structures with tailored binding interfaces. This study evaluates RFdiffusion’s efficacy in designing *de novo* binders for six targets: Strep-Tag II (a peptide tag) and five eukaryotic proteins (STAT3, FGF4, EGF, PDGF-BB, and CD4). Five binders were designed for each target and experimentally validated. While two Strep-Tag II binders outperformed streptavidin in Western blot assays, none matched the sensitivity of anti-Strep-Tag II antibodies. Binders for the other targets failed due to low expression, nonspecific binding, or undetectable affinity. Despite generating structurally diverse candidates, RFdiffusion’s success rate was limited by low-affinity designs and inconsistent recombinant expression. These results underscore the need for further optimization of AI-driven protein design tools for practical biochemical applications.

## Introduction

The development of high-affinity protein binders is a critical component in biochemical detection applications, including immunoassays, Western blotting, and mass spectrometry. These binders are essential for the accurate detection and quantification of target proteins, often serving as the foundation for diagnostic tools and research methodologies. Traditional approaches to binder design, such as antibody generation and rational design, have been successful but are often time-consuming, labor-intensive (1, 2). The advent of artificial intelligence (AI) and machine learning (ML) has introduced new possibilities for accelerating and enhancing the design of protein binders (3–6).

Among the various AI-driven methods, deep learning models have shown particular promise in the field of protein design (3). One such model is RFdiffusion (7), a RoseTTAFold-based denoising diffusion method that leverages the power of deep learning to generate protein structures with diverse topologies and optimized binding interactions. This approach has the potential to revolutionize the design of protein binders by offering a more efficient and effective alternative to traditional methods. Previous studies have highlighted the potential of AI in drug design and peptide-based drug development (8). However, the practical application of AI-generated binders in biochemical detection remains un-explored. This study aims to fill this gap by providing a comprehensive evaluation of RFdiffusion’s performance in generating binders for a range of targets. Our findings have significant implications for the future of AI in protein design and its potential to streamline and enhance biochemical detection processes.

## Results

We designed five binders targeting the Strep-Tag II peptide using RFdiffusion (an online tool from Neuronap, https://neurosnap.ai/service/RFdiffusion-v2). The input target was acquired from the PDB database with the PDB number 6QW4. The region in 6QW4 containing Strep-Tag II was selected and cropped for designing the binders. As shown in Figure 1A, the top five ranked binders were generated by RFdiffusion. Next, these binders were fused to the rabbit Fc fragment and recombinantly expressed in Expi293 cells. Two of the five binders (the 2nd and 5th) showed high expression levels in the culture supernatant, which could be detected by Coomassie-blue staining after SDS-PAGE gel separation (Figure 1B). The five culture supernatants were applied for ELISA detection, where a protein ladder (Thermo Fisher, Catalog No. 26630) containing Strep-Tag II was used as the detection target. Except for the 3rd binder, the other four binders exhibited a positive signal compared to blank wells (Figure 1C). The 2nd and 4th binders gave positive results in Western blot detection, while the 5th binder did not, despite its expression level being comparable to the 1st (Figure 1D). We purified the 2nd binder by protein A affinity chromatography, and the purity was confirmed by SDS-PAGE separation and Coomassie-blue staining (Figure 1E). The 2nd Strep-Tag II binder showed better characteristics than wild-type streptavidin, which specifically binds to Strep-Tag II and is used for recombinant protein purification (Figure 1F). However, compared to the anti-Strep-Tag II antibody, the 2nd binder was not able to detect the recombinant protein KCNQ5 and NCTP fused with the Strep-Tag II tag, possibly owing to its low affinity, which was supported by the data that the protein ladder 26630 presented a weak signal detected by the 2nd binder (Figure 1G). The differences in the 26630 bands detected by the 2nd binder between Figure 1D, Figure 1F, and Figure 1G may result from deviations in each membrane transfer during Western blotting. From these data, we demonstrated that although RFdiffusion could generate protein binders with expected functions to some extent, the efficacy of the binders was still not as high as that of antibodies.

**Figure 1.**
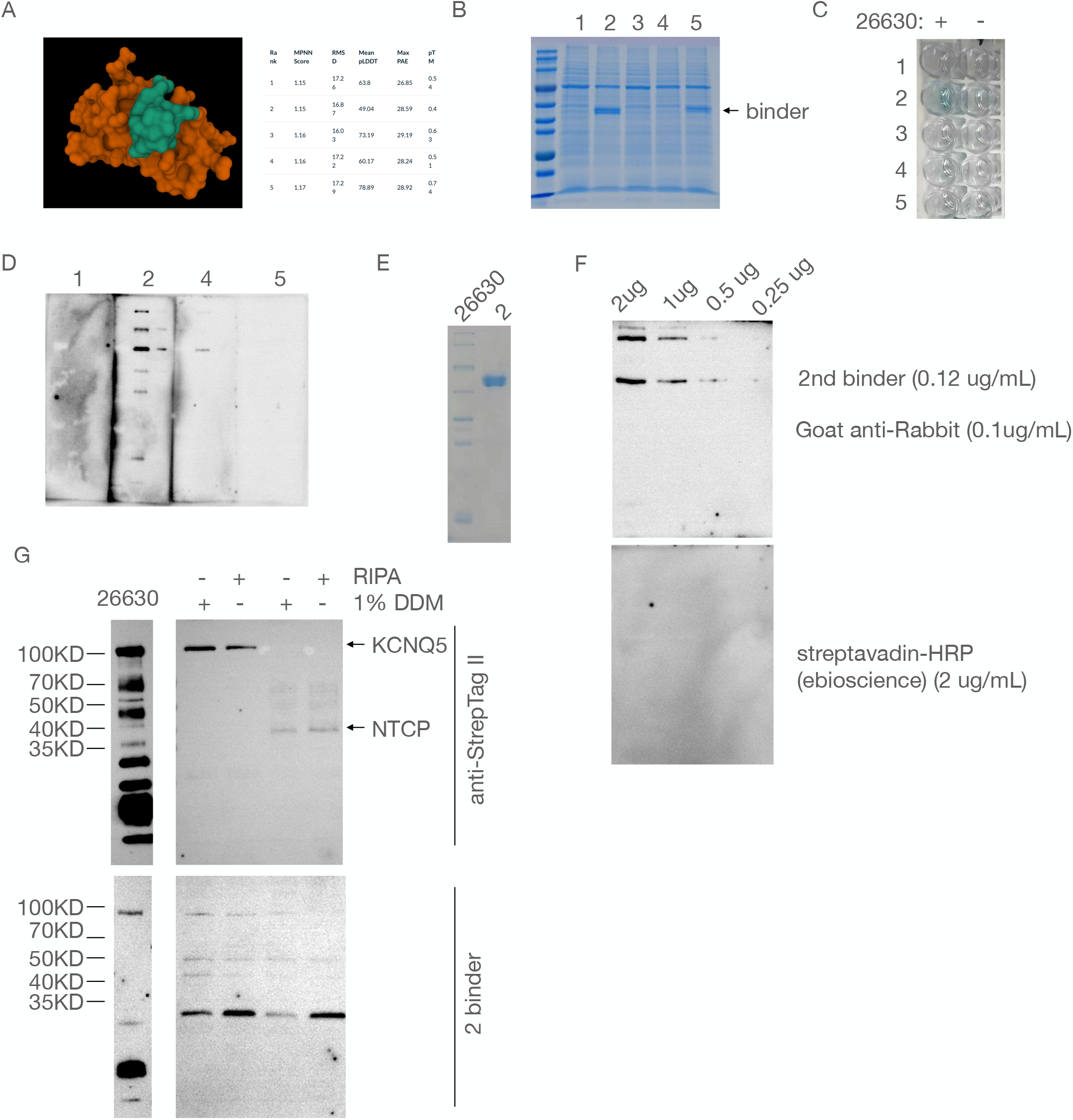
Binders against short peptide StrepTagII is disadvantageous to anti-StrepTagII antibody. (A) Five binders were designed by RFdiffusion. The critical parameters were shown. A online website Neuronap that using RFdiffusion was used. (B) The binders fused with mouse Fc tag were expressed in Expi293 cells. The culture supernatant was analysed by SDS-PAGE separation. (C) Elisa confirmation of binders capacity binding to StrepTagII. The protein ladder from thermo fisher with the Catalogue number 26630 was used as the coating antigen. Of note, 26630 contained the proteins tagged with StrepTagII. (D) Western-blot analysis of binders binding to 26630. (E) The 2nd binder was purified by proteinA column and purity was confirmed by SDS-PAGE separation. (F) Comparison of 2nd binder and streptavadin for western-blot analysis. 26630 was loaded in a serial dilution. (G) Comparison of 2nd binder and anti-StrepTag II antibody for western-blot analysis. HEK293 cells expressing two transmembrane protein NTCP and KCNQ5 was used as the detection sample.

Besides the peptide target, we further tested five protein targets for de novo binder design. Briefly, the targets used for binder binding were acquired from the PDB database, and the region for binder contact was further cropped or a hotspot site was selected before RFdiffusion design. Five binder candidates targeting STAT3 (fused with rabbit Fc) were chosen, but only one binder showed detectable expression from Expi293 culture supernatant (data not shown). Compared to the anti-STAT3 antibody, no specific bands were detected by the STAT3 binder (Figure 2A). A similar result was shown by immunofluorescence detection (Figure 2B). Furthermore, we chose CD4, a surface marker of CD4+ T cells, for binder design. Only one binder against CD4 could reach a detectable expression level. However, no binding capacity was detected by flow cytometry (data not shown).

**Figure 2.**
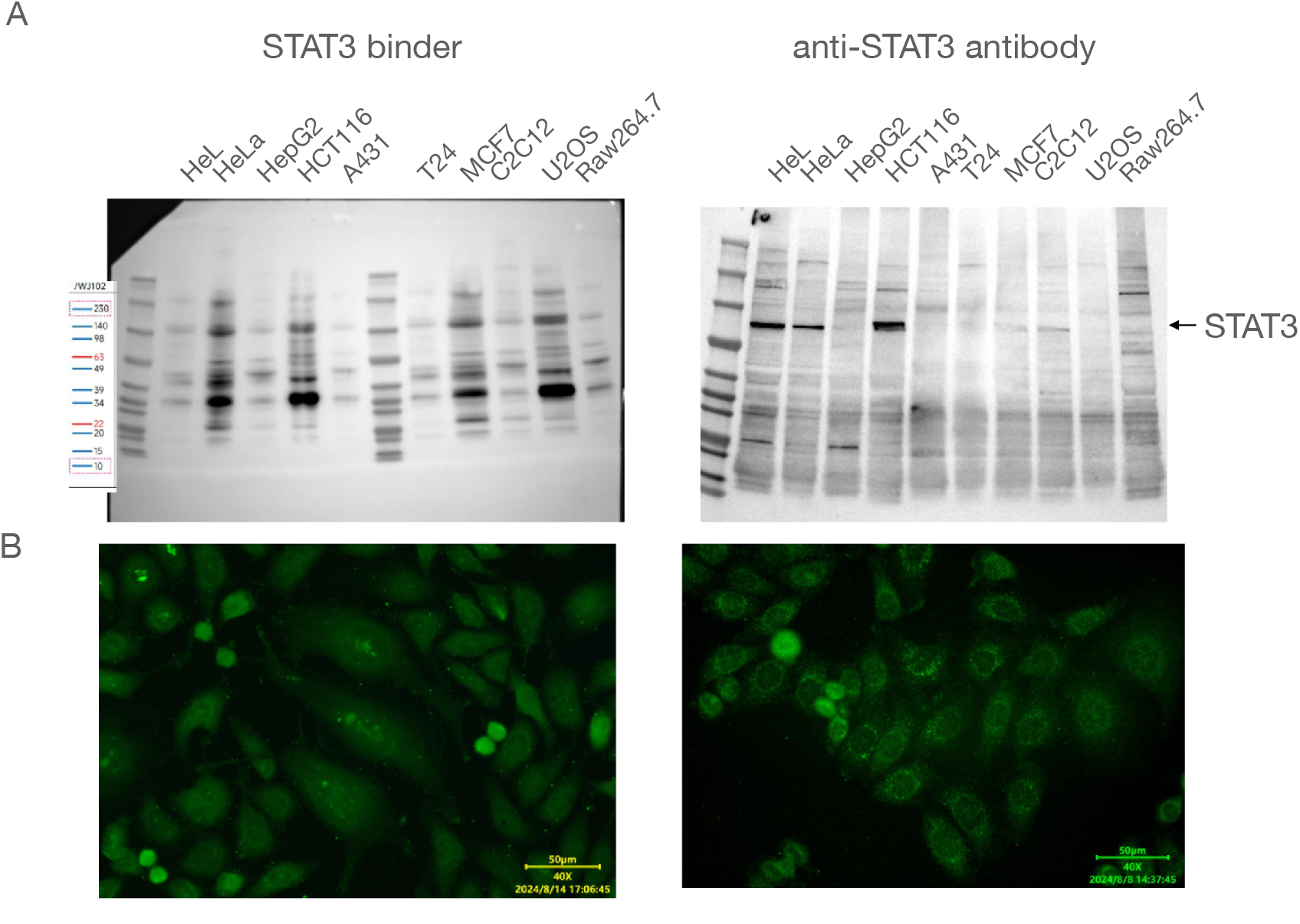
Binders against STAT3 is not able to apply to western-blot or immunofluoscence detection. (A) Western blot analysis of STAT3 expression in different cell lines. Cell protein sample with 10ug by RIPA preparation was loaded for SDS-PAGE separation, and further transferred into the NC membrane. Then, binder or the primary antibody was incubated 1h at RT and washed 4 times with TBST. The secondary antibody HRP conjugated goat anti-mouse was incubated 1h at RT. (B) HeLa cells was used for immunofluoscence analysis. 4%PFA was used for fixation 5 min at RT and 0.05% TritonX100 was used to treat the cells 5 min at RT. Then, binder and antibody was incubated and washed 3 time by PBS, followed by FITC-conjugated goat anti-mouse. Photos were captured by the Olympus IX53.

Finally, we designed binders targeting three cytokines: FGF4, EGF, and PDGF-BB. Instead of fusing with Fc, these 15 binders were fused with NusA and expressed in E. coli BL21 (Figure 3A). No strong binding was observed between the targets and binders (Figure 3B, 3C). Together, these data demonstrated that RFdiffusion was able to design new binders but did not exhibit a high success rate. The reason for this low success rate may result from the limited number of binder candidates (five binders for each target), the low level of recombinant expression, and the low affinity due to the algorithm in RFdiffusion. Our results indicated that RFdiffusion may not substitute for conventional binder development methods, such as antibody generation.

**Figure 3.**
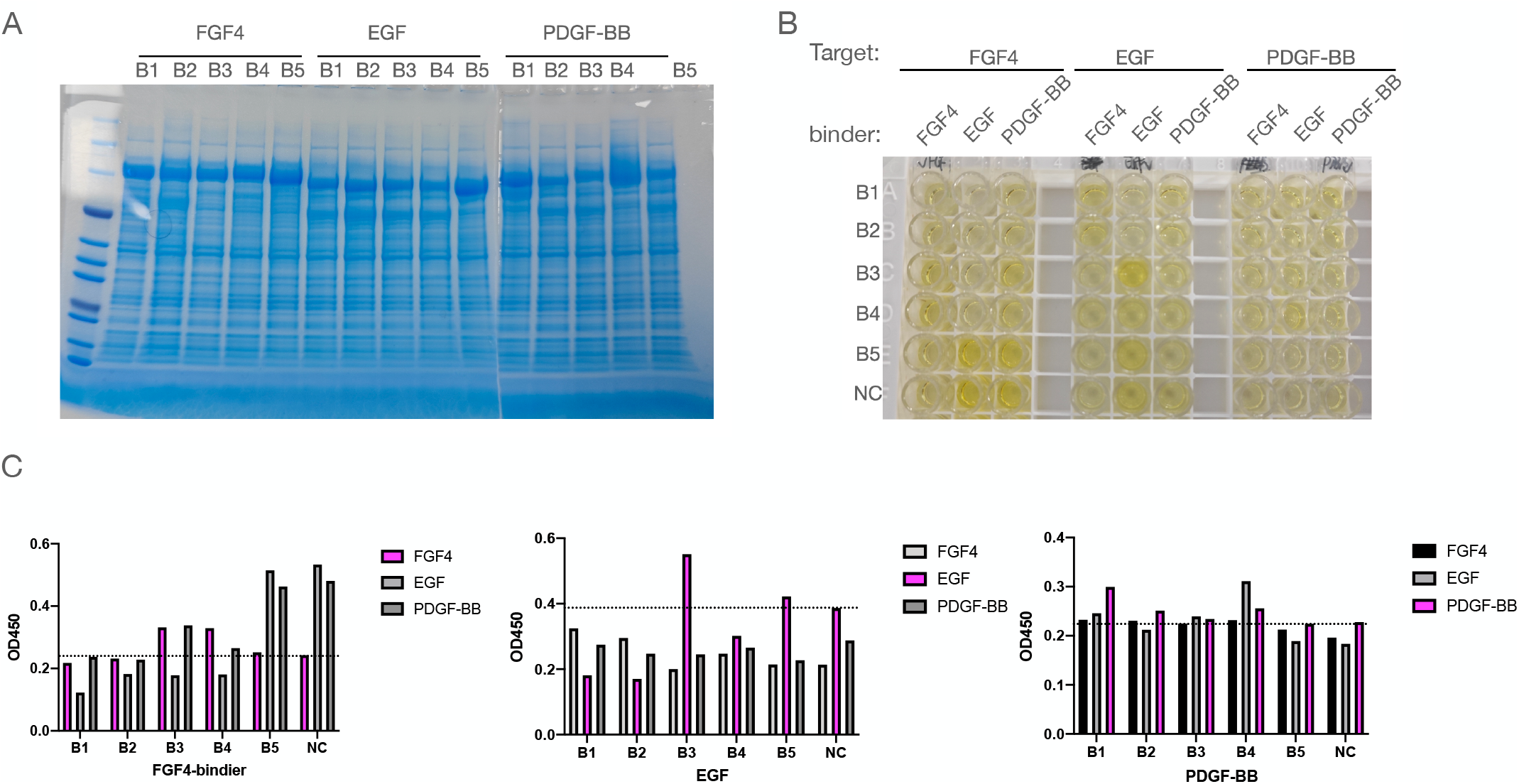
Binders against cytokines FGF4, EGF or PDGF-BB are not able to apply to Elisa detection. (A) The binders fused with NusA tag were expressed in E-coli BL21 cells. The culture supernatant was analysed by SDS-PAGE separation. (B) Elisa detection of binders capacity of binding to cytokines. Recombinant cytokines were coated in the Elisa plate for detection. (C) Statistic of OD 450 in B.

